# A neuroimmune IL-13 axis is associated with human enteric nervous system development

**DOI:** 10.64898/2026.05.27.728110

**Authors:** Luís Eduardo Duarte Gonçalves, Bruno Ghirotto, Alejandra Pizano, Lotta Roßdeutsch, Oana-Maria Thoma, Anne Jacobsen, Leah Trumet, Manuel Weber, Stephan P. Rosshart, Manuel Besendörfer, Markus Eckstein, Carolin Schmidt, Atefeh Namipashaki, Hanchen Yu, Marlene M. Hao, Lincon A. Stamp, Markus Neurath, Claudia Günther, Beate Winner, Inga V. Hensel, Sonja Diez

## Abstract

The mechanisms governing maturation of the human enteric nervous system (ENS) during early postnatal life remain poorly defined. Here, we characterize the transcriptomic, cellular, and functional landscape of the neonatal human ileum and identify a neuro-immune axis associated with ENS expansion. Using human tissue transcriptomics, flow cytometry, iPSC-derived enteric neural lineages, and single-cell interactome analyses, we show that the neonatal ileum is enriched for pro-neurogenic transcriptional programs and harbors a greater abundance of enteric neurons and glia than the adult tissue. T cells emerge as a predominant source of interleukin-13 (IL-13) in the neonatal gut, and enteric neurons express its receptor IL13RA1, enabling direct immune-to-neuron signaling. Functional experiments demonstrate that IL-13 enhances expression of key enteric neuronal markers in a concentration-dependent manner. In parallel, single-cell analyses identify enteric neurons as a major predicted source of macrophage migration inhibitory factor (MIF), with signaling directed toward T and NK cell populations, suggesting that the ENS actively shapes the immune environment it depends upon. Together, these findings support a model in which bidirectional neuro-immune communication establishes a pro-neurogenic niche during a critical window of ENS development. This work positions the neonatal immune system as an active contributor to ENS maturation and offers a new perspective on how neuro-immune crosstalk shapes intestinal development in early life.

## Introduction

The gastrointestinal tract is innervated by the enteric nervous system (ENS), an intrinsic neural network composed of interconnected neurons and glia organized into myenteric and submucosal plexuses. Positioned at the interface of the epithelial barrier and mucosal immune compartment, the ENS occupies a central role in coordinating host–microbiome interactions and neuro-immune communication.

The early postnatal period represents a critical window of intestinal adaptation, during which the newborn transitions from placental nutrition to enteral feeding and undergoes rapid microbial colonization (1). This transition is accompanied by extensive remodeling of the epithelial and immune compartments, including the establishment of barrier integrity and the maturation of immune responses. Although the ENS has traditionally been viewed primarily as a regulator of gastrointestinal motility and secretion, emerging evidence highlights its broader role as an active participant in neuro-immune communication (2). While neonatal-adult differences in the epithelial and immune compartments are increasingly well characterized (3, 4), the developmental trajectory of the ENS during early life remains comparatively poorly understood. Although, basic ENS architecture is largely established at birth, substantial maturation is required before the system to acquire the sensory and effector capabilities of the adult intestine, and the signals governing this process have not been defined.

Recent advances in molecular profiling have enabled detailed characterization of the ENS, revealing pronounced age- and species-dependent differences in cellular composition and function (5). Beyond its classical roles, the ENS regulates immune and epithelial cell function through the secretion of cytokines, neurotransmitters, and growth factors (6-8). ENS-derived mediators such as interleukin-18, vasoactive intestinal peptide (VIP), and acetylcholine contribute to epithelial barrier maintenance by promoting antimicrobial peptide production and mucus secretion (9-12). The importance of the neuro-immune interface is underscored in pathological conditions such as Hirschsprung disease, where aganglionic segments of the colon exhibit not only impaired motility but also disrupted mucosal immunity and altered goblet cell function (13, 14).

The ENS is itself responsive to immune-derived signals. Under homeostatic and inflammatory conditions, epithelial and immune cells release mediators that directly modulate neuronal activity and function. Notably, type 2 cytokines including interleukin-5 and interleukin-13 influence neuronal signaling programs and are sufficient to induce neurons to release immunomodulatory neuropeptides (15), highlighting the bidirectional nature of neuro-immune communication in the gut. However, knowledge on neuro-immune communication shaping ENS development remains sparse.

A fundamental gap remains in understanding how the ENS develops during early life, and in particular, how interactions with the immune system shape this postnatal maturation. Here, we address this gap by characterizing the transcriptomic, cellular, and functional landscape of the neonatal human ileum, and identify a neuro-immune axis in which T cell-derived IL-13 and neuronal MIF signaling converge to establish a pro-neurogenic environment during a critical window of ENS development. These findings offer a new perspective on the establishment of gut function in early life and on the origins of gastrointestinal disorders in which neuroimmune homeostasis is disrupted.

## Results

### Neonatal Ileum Is Enriched for Enteric Neurons

To identify age-dependent cues in the enteric nervous system, we characterized the transcriptomic landscape of neonatal and adult ileal full-thickness resections (Fig. 1A, Table 1). Ileal tissue was collected from six neonates without any connection to inflammation (atresia, ileus), all being fed with breast milk and representing balanced preterm to term gestational ages, and four adults with no inflammatory gastrointestinal diseases. Given the clinical vulnerability of the neonatal ileum to neuro-inflammatory conditions like NEC and HSCR we decided to focus our analysis on this intestinal segment. Bulk RNA-sequencing revealed pronounced differences between the developing and mature intestinal tissue. In neonates, innate immune pathways were preferentially activated, including interleukin-1 receptor binding, neutrophil migration, and antimicrobial peptide production. In contrast, the adult gut was characterized by a shift toward adaptive immunity, with enrichment of Th1/Th2 and Th17 cell differentiation programs and an intestinal IgA production network. Published results by others suggested a potential infiltration of adaptive immune cells during the first 3 months of life (Fig. 1B) (16). Strikingly, neonatal samples also exhibited significant enrichment of neurodevelopmental gene sets, including NCAM signaling for neurite outgrowth, axon development, enteric nervous system development, neurogenesis, and neuron migration (Fig. 1C), suggesting active ENS maturation during this early postnatal window.

**Fig. 1.**
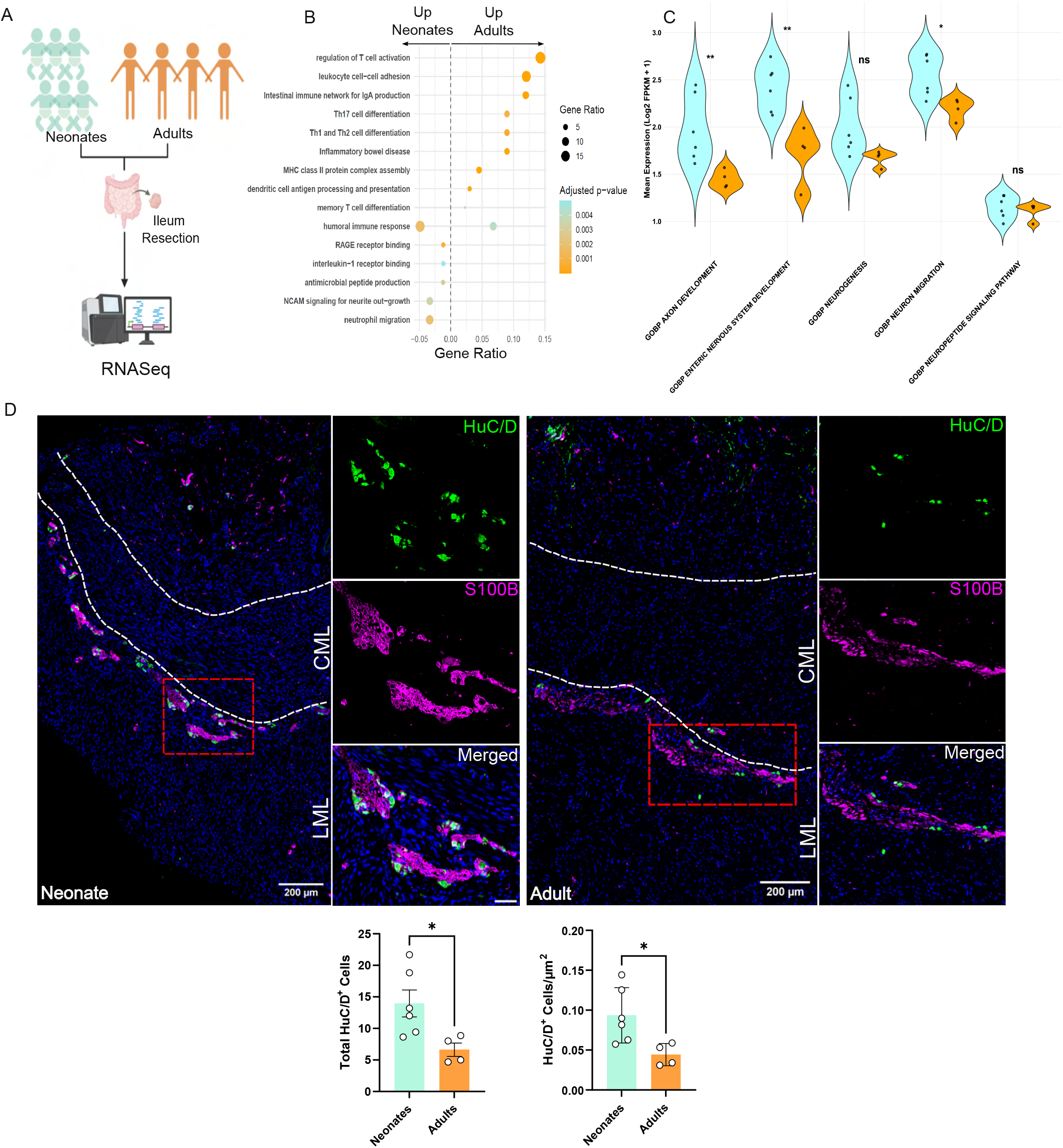
Characterization of Enteric Nervous System Landscape in Neonates. (A) graphical representation (created with BioRender) illustrating the collection of ileum resections for bulk RNA-Sequencing. (B). Dot plot displaying differentially regulated pathways in neonates versus adults. Significance was defined by an adjusted p-value (padj < 0.05). (C) Violin plots representing the mean normalized expression of genes within neuronal-related pathways. Statistical significance was determined using the Wilcoxon Rank-Sum test (* p < 0.05, ** p < 0.01). (D) Representative images of neonatal and adult ileum tissues stained for neuronal (HuC/D) and glial (S100B) markers. Quantification displays the density of HuC/D+ nuclei normalized per um2. CML= circular muscular layer; LML=longitudinal muscle layer. Scale bar = 10µm. For all panels, n=6 neonates and n=4 adults

**Table 1.** Patients’ demographics. Clinical characteristics of the 10 included patients of the study.

| Characteristics | Neonates (n=6) | Adults (n=4) |
| --- | --- | --- |
| **Sex [n]** |  |  |
| Male | 3 | 1 |
| Female | 3 | 3 |
| **Age at collection of tissue [median (range)]** | 3 months (2–6 months) | 65 years (25–88 years) |
| **Gestational age [median in weeks (range)]** | 26 (23–37) | – |
| **Weight at birth [median in grams (range)]** | 625 (580–3580) | – |
| **Diagnosis [n]** |  |  |
| Meconium Ileus | 4 | 0 |
| Intestinal stenosis | 2 | 0 |
| Colon/Rectal Carcinoma | 0 | 3 |
| FAP | 0 | 1 |
| **Surgical indication due to [n]** |  |  |
| Ostomy removal | 5 | 1 |
| Intestinal stenosis | 1 | 0 |
| Colectomy | 0 | 3 |

To validate these findings at the protein level, and add a spatial resolution, we performed immunostaining of neonatal and adult ileum samples using established markers for enteric neurons (HuC/D) and glia (S100B) (17). Both cell types were organized into ganglionic structures within the mucosal and submucosal layers, where neurons and glial cells were found in direct contact (Fig. 1D). Quantification revealed an approximate two-fold increase of HuC/D+ neurons within ganglia of the neonatal compared to adult ileum (Fig. 1D). Taken together, transcriptomics and histological data indicate that the neonatal ileum is a site of active enteric neuronal expansion and maturation.

### IL-13 as a Candidate Mediator of Neonatal Enteric Neurogenesis

Given the pronounced enrichment of neurodevelopmental pathways in the neonatal ileum, we sought to identify key molecular signals that might drive this age-specific program. Among the differentially expressed cytokines, interleukin-13 (IL-13) emerged as a compelling candidate (Fig. 2A). IL-13 is a central regulator of gut immunity, known to modulate neurotransmitter production—such as neuromedin U (NMU) and calcitonin gene-related peptide (CGRP)—and to fine-tune immune responses to helminthic infection (15). Beyond its immune functions, IL-13 has been shown to promote synaptic plasticity, neurogenesis and protection in the central nervous system (18). These pleiotropic roles suggest a potential role in peripheral nervous system development. We therefore hypothesized that IL-13 may serve as a critical link between early life immune activation and enteric neuronal maturation in the neonatal gut. To test this, we first assessed the expression of IL-13 receptor subunit IL13RA1 in our cohort. IL13RA1 was markedly upregulated in neonatal samples compared to adults (Fig. 2A). However, since IL13RA1 is not ubiquitously expressed across the ENS and bulk RNA-seq lacks cellular resolution (5, 19), we re-analyzed single-cell RNA sequencing (scRNA-seq) data from an independent cohort of neonatal gut tissue (aged 2-5 months) (17). After subsetting and UMAP clustering, we identified seven distinct enteric neuron subclusters (Fig. 2B). Notably, the vast majority of these subclusters expressed high levels of IL13RA1, while IL13RA2 expression was minimal or absent (Fig. 2C). This indicates that enteric neurons in the neonatal ileum are equipped to respond to IL-13 signaling via IL13RA1. Our data showed that both IL-13 and IL13RA1 were significantly enriched in neonatal compared to adult samples, in contrast to other immune pathways such as Th1 (IFNG) and Th17 (RORC), which showed less consistent expression patterns. (Fig. 2A). To confirm receptor presence at the protein level, we performed immunofluorescence of IL13RA1 and HuC/D in enteric glia from neonatal and adult ileum (Fig. 2C). Receptor expression was observed in neonatal ganglia, with weaker but detectable signals in adults. Together, these data demonstrate that enteric neurons in the neonatal ileum express functional IL-13 signaling machinery, suggesting a potential role for IL-13 in shaping ENS development during early postnatal life. This finding positions IL-13 as a candidate mediator linking immune activation and neurodevelopment in the maturing gut.

**Fig. 2.**
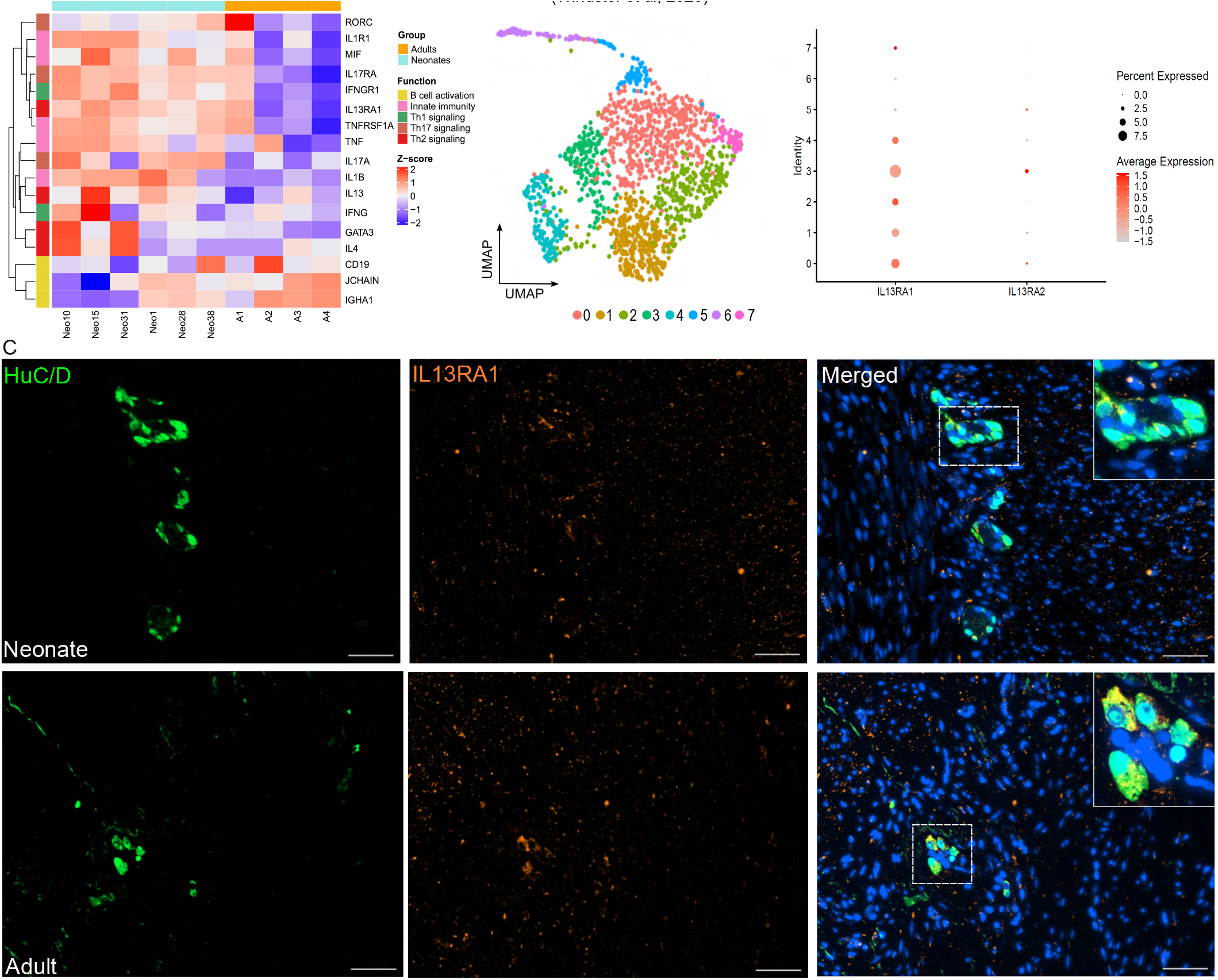
Expression and Localization of IL-13RA1 in the Neonatal and Adult Ileum. (A) Heatmap illustrating the transcriptomic profiles of diverse immune compartments within the ileal mucosa. (B) UMAP embedding of enteric neuronal subclusters identified in the ileum (left), with corresponding feature plots (right) mapping the specific expression patterns of the IL13RA1 receptor across neuronal subtypes. (C) Representative immunofluorescence micrographs confirming the protein-level expression of IL-13RA1; white arrows indicate colocalization of the receptor within HuC/D+ neurons in the myenteric plexus of the ileum. Scale bar = 10µm. For all panels, n=6 neonates and n=4 adults

### T Cells are the Primary Source of IL-13 in the Neonatal Ileum

Given the prominent role of IL-13 in enteric neuron-mediated immunity and gut homeostasis, we next aimed to identify the cellular source of IL-13 in the neonatal and adult ileum. To Simultaneously quantify neuronal and leukocyte populations, we performed flow cytometry using an established ENS gating strategy (17) complemented by a panel of immune cell markers (Fig. 3A – S1A-B).

**Fig. 3.**
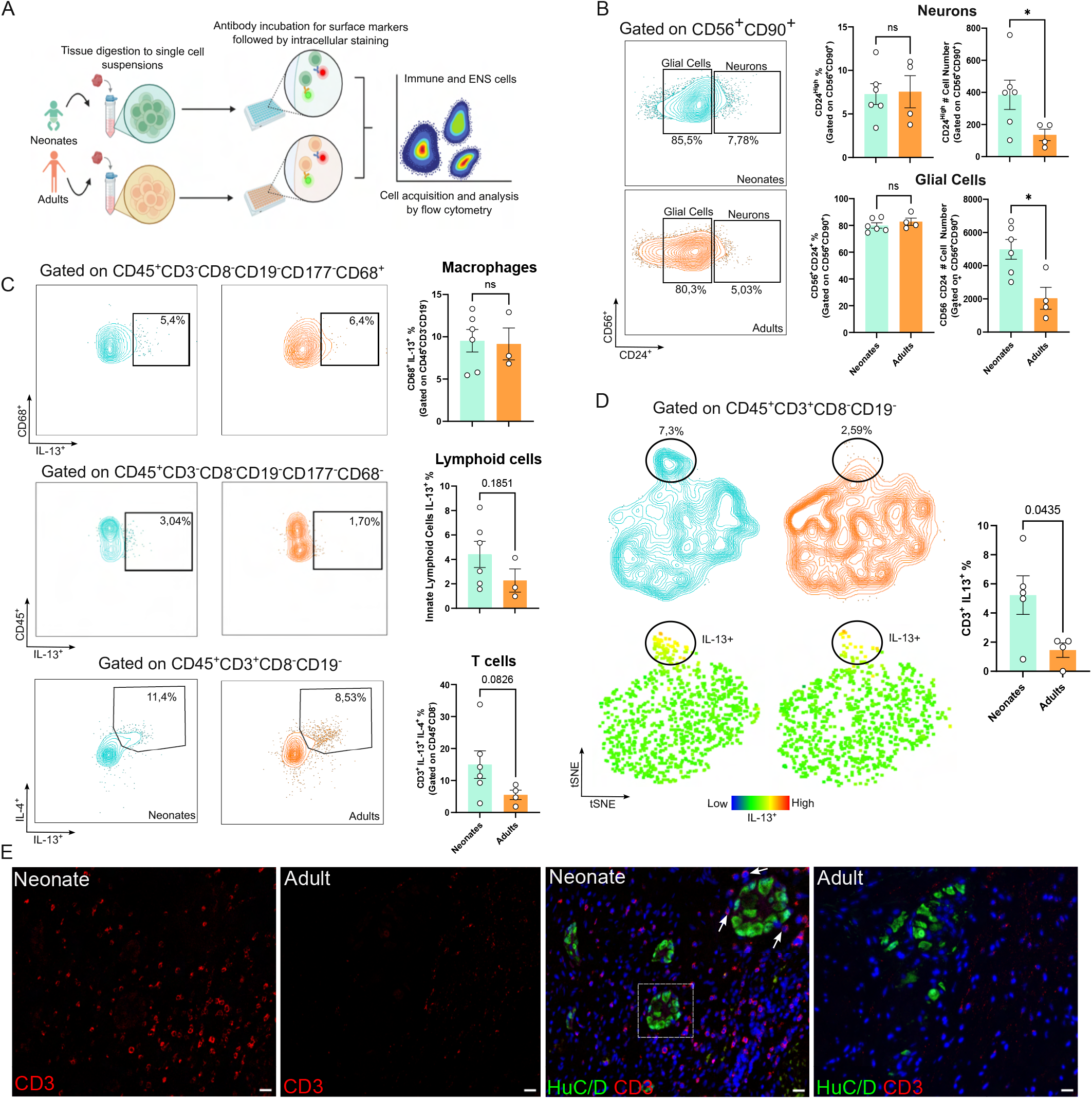
Characterization of IL-13 Expression Across Immune and Enteric Nervous System Compartments. (A) Schematic workflow illustrating the experimental pipeline for ileal tissue digestion and the generation of single-cell suspensions for downstream analysis. (B) Representative flow cytometry gating strategy for the enteric nervous system, with neurons identified as CD56+CD90+CD24high and glial cells defined as CD56+CD24+. Accompanying bar graphs provide the absolute quantification of neuronal and glial populations in both neonatal and adult cohorts. (C) Frequency of intracellular IL-13 expression across key neonatal immune subsets, including macrophages, innate lymphoid cells (ILCs), and T cells (* p < 0.05). (D) t-SNE visualization of the T cell compartment highlighting the distribution and relative intensity of IL-13 expression within T cell subpopulations in neonates compared to adults. (E) Immunofluorescence micrographs of T cells (CD3+) in proximity to neurons (HuC/D+) in neonatal and adult ileal sections. Scale bar = 10µm. For all panels, n=6 neonates and n=4 adults

We used CD90, CD56 and CD24 to characterize the ENS compartment. CD90 (Thy-1) is a glycosylphosphatidyli-nositol (GPI)-anchored surface glycoprotein commonly expressed on mesenchymal and neuronal cells and is widely used to delineate stromal and neuro-associated cell populations. CD56 (neural cell adhesion molecule, NCAM) functions as a key mediator of cell–cell adhesion in neural tissues, while CD24 is a small GPI-anchored surface protein typically associated with immature or progenitor-like phenotypes, enabling further phenotypic stratification of the ENS compartment.

This quantification confirmed a significantly higher number of enteric neurons and glia in neonates compared to adults (Fig. 3B), consistent with our transcriptomic and histological findings (Figure 1B-D). We then examined candidate IL-13-producing populations including T-helper 2 (Th2) cells, type 2 innate lymphoid cells (ILC2s), and macrophages (19). Intracellular IL-13 levels were unchanged in ILCs and macrophages between neonatal and adult ileum (Fig. 3C), whereas T cells showed a trend to increased IL-4/IL-13 production in neonates. Unsupervised tSNE analysis further revealed that IL-13-producing T cells reside within a phenotypically distinct neonatal cluster (Fig. 3D). Next, we asked whether T cells were spatially positioned to communicate with enteric neurons. We examined ileal sections by immunostaining. CD3+ T cells were enriched in the immediate vicinity of HuC/D+ ganglionic clusters in neonatal but not adult tissue (Fig. 3E). Together, these findings identify T cells as the primary source of IL-13 in neonatal ileum, raising the possibility that T cell-derived IL-13 acts on neuronal IL-13 receptors expressed in enteric ganglia.

### IL-13 enhances expression of enteric neuron markers in iPSC-derived enteric neural lineages

Having identified T cells as the primary source of IL-13 (Fig. 3) and enteric neurons expressing its receptor IL13RA1 (Fig. 2), we next asked whether IL-13 can directly drive neurogenic programs, given its reported association with neurodevelopment (20). We first examined this relationship in our bulk RNA-seq data from human ileal biopsies. IL13RA1 expression showed a positive correlation trend with the enteric neuronal marker ELAVL4 (HuC/D) in neonates (r = 0.78, p = 0.065; Fig. 4A), with a more pronounced association than in adults (r = 0.66, p = 0.34), suggesting that IL-13 signaling may be linked to factors regulating ELAVL4 expression during early postnatal development. To functionally test whether IL-13 influences enteric neurogenesis, we used a human induced pluripotent stem cell (iPSC)-derived enteric neural lineage model generating mixed populations of neurons and glial cells (21) (Fig. 4B).

**Fig. 4.**
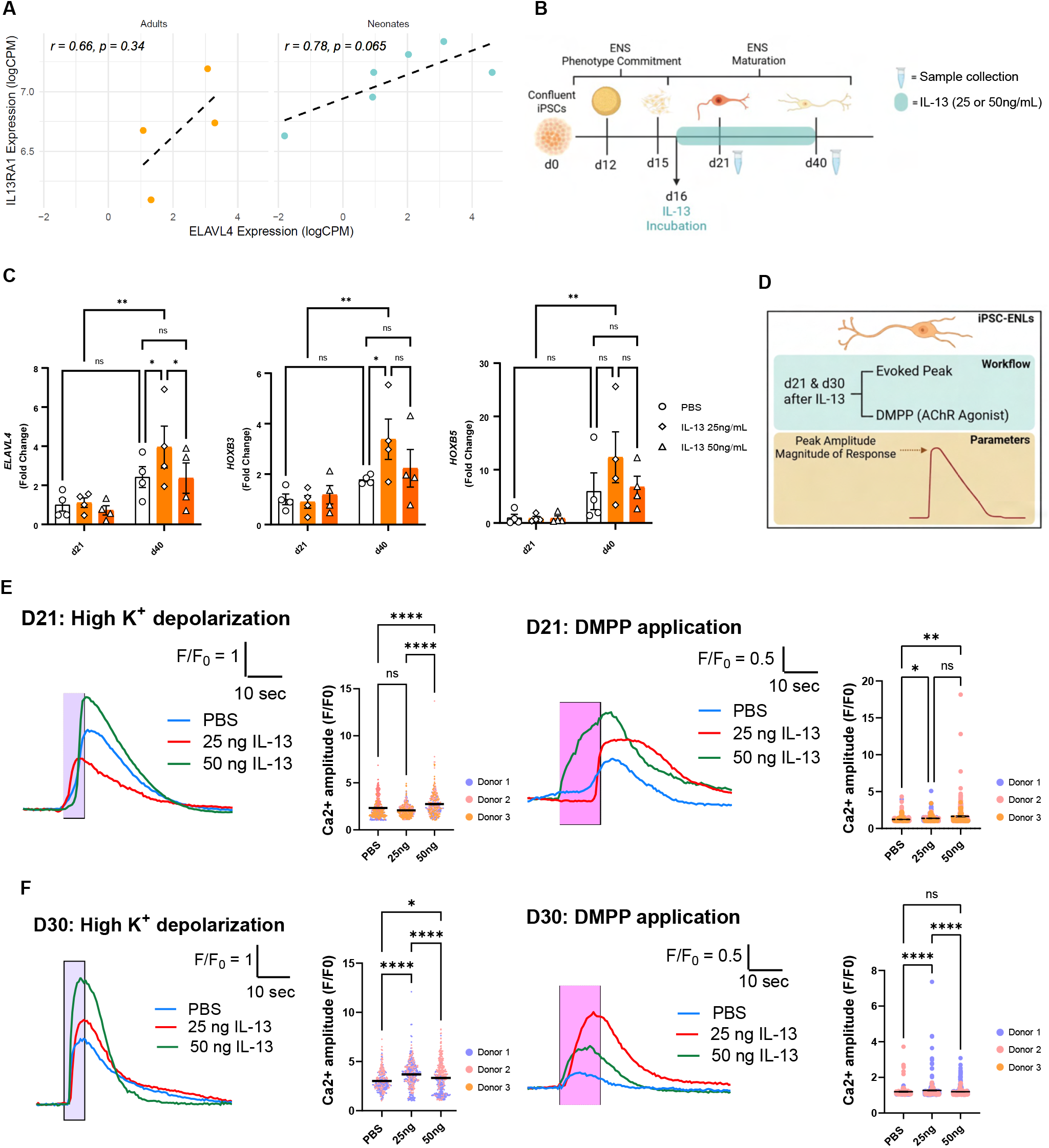
IL-13 activates pro-neurogenic machinery in iPSC-derived ENS. (A) Correlation analysis of IL13RA1 and ELAVL4 (encoding HuC/D+) expression levels in neonatal and adult ileal tissues, highlighting the relationship between receptor presence and neuronal density, n=6 neonates and n=4 adults. (B) Experimental schematic (created with BioRender) depicting the differentiation of human induced pluripotent stem cells (iPSCs) into enteric neurons and glia, followed by recombinant IL-13 stimulation at concentrations of 25 ng/mL or 50 ng/mL. (C) Quantitative expression analysis of key pro-neurogenic and pan-neuronal markers (ELAVL4, HOXB3, HOXB5, ASCL1, and TUBB3) following IL-13 treatment at early (Day 21) and late (Day 40) stages of differentiation, n=4 independent donors. (D) Experimental paradigm for single cell calcium imaging experiments. Generated with Biorender. (E) Functional maturation at d21. Quantification of Ca2+ responses to High K+ (left, n=425, 505 and 323 cells for PBS, 25ng/mL and 50ng/mL IL-13, respectively, from three independent biological donors) and DMPP (right, n=319, 168 and 283 cells for PBS, 25ng/mL and 50ng/mL IL-13, respectively, from three independent biological donors) in iPSC-derived enteric neural lineages (iPSC-ENLs) treated with PBS, 25ng/mL, or 50ng/mL IL-13. (F) Functional maturation at d30. Quantification of Ca2+ responses to High K+ (left, n=276, 351 and 423 cells for PBS, 25ng/mL and 50ng/mL IL-13, respectively, from two independent biological donors) and DMPP (right, n=123, 277 and 362 cells for PBS, 25ng/mL and 50ng/mL IL-13, respectively, from two independent biological donors).

Enteric neural crest progenitors from four healthy donors were treated with IL-13 at 25 or 50 ng/mL during the ENS maturation phase (day 16-40), to mimic early neonatal conditions (Fig. 4B). Expression analysis at day 40 (d40) revealed that the lower dose (25ng/mL) significantly upregulated key enteric neuronal markers, ELAVL4, HOXB3 and HOXB5, whereas the higher dose (50ng/mL) produced no significant effect to controls, indicating that IL-13-driven neurogenic activity is concentration-dependent (Fig. 4C).

To evaluate the functional impact of IL-13 on enteric lineage maturation, we performed live cell calcium imaging at two developmental stages, d21 and d30, in response to high K+ depolarization and the nicotinic receptor agonist, 1,1-dimethyl-4-phenylpiperazinium iodide (DMPP) (Fig. 4D-F). Changes in intracellular calcium were monitored across individual cells, and the magnitude of responses showed a clear dose-dependent maturation. During early neuronal development (d21), increasing dose of IL-13 treatment had greater impacts on Ca2+ responses, with 50 ng/mL treatment groups showing higher depolarization-evoked [Ca2+] amplitudes (Fig. 4E). Responses to DMPP were increased following treatment with both 25 ng/mL and 50 ng/mL IL-13 (Fig. 4E). As the lineages matured to d30, treatment with 25 ng/mL showed significantly higher responses to both high K+ depolarization and DMPP (Fig 4F). Together, these data indicate that while high-dose IL-13 may influence early calcium handling kinetics, the near-physiological 25 ng/mL dose is sufficient to drive a sustained increase in total neuronal responsiveness and cholinergic sensitivity.

### MIF Signaling Links Enteric Neurons to T Cells in the Neonatal Ileum

To validate our findings in an independent cohort and define the signaling architecture of the neonatal ileum at single-cell resolution, we analyzed integrated scRNA-seq data from the Gut Cell Atlas (22) (Fig. 5 and Fig. S2), comprising 25 datasets of ileal samples from healthy neonates and adults. UMAP analysis revealed distinct cellular compartments, including a prominent neural cluster, immune populations of B, T, myeloid, and NK cells, as well as epithelial and mesenchymal stromal populations (Fig. 5A). The neural compartment was significantly enriched in the neonatal ileum, while T, NK, and plasma cell populations were reduced relative to adults (Fig. 5A–B and Fig. S2A–B), consistent with our primary observations. To characterize intercellular communication, we performed cell-cell interaction analysis using CellChat (23). While the overall number and strength of predicted interactions were comparable between neonatal and adult samples, the macrophage migration inhibitory factor (MIF) signaling pathway emerged as a prominent feature of the neonatal ileum (Fig. 5C–E and Fig. S2C–E). Enteric neurons were identified a major predicted source of MIF, with inferred interactions targeting CD74, CXCR4, and CD44 on T and NK cells (Fig. 5C–D and Fig. S2E–G), and neuronal MIF signaling was elevated in neonates compared to adults (Fig. S2E–G). Accordingly, our own RNA-seq data highlighted the higher expression of MIF in the neonatal ileum (Fig. 2A). Network analysis additionally identified FCER2A (CD23) signaling in neonatal B cells, associated with Th2 response development (24), and retinoic acid signaling within the neural compartment, a pathway essential for enteric neuron development (25) (Fig. 5C). Examining cytokine expression within immune populations, neonatal T cells in this independent cohort showed significantly higher expression of IL-13 and IL-4 expression than adult T cells (Fig. 5E), in line with our flow cytometry data (Fig. 3D). Downstream signaling mediators associated with IL-13 signaling, including STAT3 and STAT6, were correspondingly elevated in neonatal neural populations (Fig. 5F), indicating active cytokine signaling within the neonatal ENS.

**Fig. 5.**
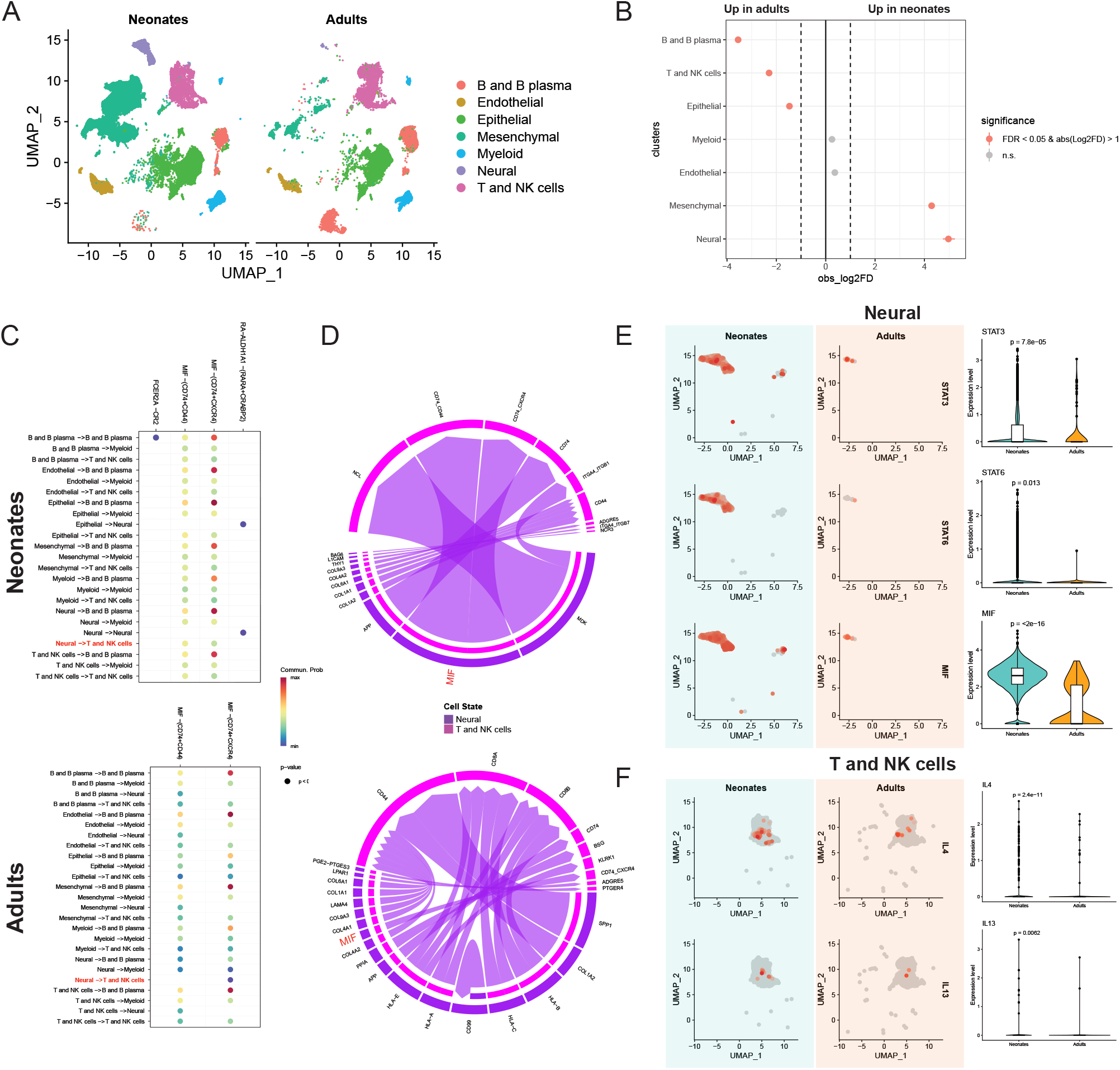
MIF Acts as a Potential Mediator Between T Cells and Enteric Neurons in the Ileum. (A) UMAP embedding illustrating the diverse neural and immune cell populations identified within the neonatal and adult small intestine. (B) Comparative statistical analysis quantifying the relative abundance of each identified cell population between neonatal and adult cohorts. (C) Ligand-receptor communication network analysis mapping the global signaling interactome across all small intestinal cell types, highlighting specific mediators and their respective target populations. (D) Chord plot visualizing the enriched MIF-mediated signaling axis, specifically depicting increased communication from enteric neurons toward T cell and natural killer (NK) cell populations. (E) UMAP plots showing expression of STAT3, STAT6 and MIF in the neural population, followed by a violin-plot quantification using the Wilcoxon test. (F) UMAP plots showing expression of IL4 and IL13 in the T and NK cell population, followed by a violin-plot quantification using the Wilcoxon test.

Together, these single-cell analyses support a model of coordinated neuro-immune signaling in the neonatal ileum, in which neuronal MIF expression, Th2-derived cytokines, and downstream neuronal signaling converge to establish a proneurogenic environment during early postnatal development.

## Discussion

The postnatal transition represents a profound physiological turning point during which the ENS must rapidly adapt to enteral nutrition and microbial colonization. Intestinal ENS density per mm2 is higher in neonates compared to adults in mice and humans. This is largely due to the massive elongation and circumferential expansion of the gut tube postnatally, effectively stretching out the space between ganglia (26-28). While fetal studies have provided further critical insights into ENS structural maturation (29, 30), our findings identify the neonatal period as a distinct phase of active ENS expansion. The neonatal ileum is characterized by a pronounced pro-neurogenic transcriptional landscape and an increased abundance of enteric neurons and glial cells relative to adult tissue, demonstrating that ENS maturation continues well beyond birth and underscoring the limitations of animal models in capturing human-specific postnatal developmental trajectories (31).

Our data support a neuro-immune axis underlying this early life ENS expansion, in which the neonatal immune compartment functions as a developmental scaffold rather than solely a mediator of host defense. T cell–derived IL-13 emerges as a key component of this interaction. Although classically associated with type 2 inflammation, IL-13 acts here in a fundamentally different capacity: enteric neurons prominently express IL13RA1, providing them with the capacity to directly integrate immune-derived signals. Functional validation using human iPSC-derived enteric neural lineages confirms that IL-13 is sufficient to enhance expression of key neuronal markers at physiologically relevant concentrations. This was further confirmed by calcium imaging, where 25 ng/mL IL-13 significantly boosted evoked Ca2+ amplitudes and enhanced sensitivity to cholinergic stimulation, despite high-dose IL-13 initially perturbing early firing kinetics. These findings align with emerging evidence that IL-13 modulates neural function in both the enteric and central nervous systems (15, 18, 32). Critically, the neurogenic effect is concentration-dependent, suggesting that under homeostatic conditions IL-13 acts as a neurotrophic factor rather than an inflammatory mediator, consistent with the non-pathological densities of T cells observed surrounding enteric ganglia in neonatal tissue. Our integrative scRNA-seq analysis suggests MIF as an upstream regulator of this neuro-immune crosstalk. Enteric neurons emerge as a major predicted source of MIF in the neonatal ileum, which was also upregulated in this group in our RNA-seq analysis, with signaling directed toward immune populations expressing CD74, CXCR4, and CD44. Given MIF’s established role as a chemoattractant, neuronal-derived MIF may actively recruit or retain the IL-13–producing T cells that sustain the neurogenic niche. The additional enrichment of Th2-associated signaling pathways like FCER2A (CD23) (24) and retinoic acid signaling, a known driver of enteric neurogenesis (25), suggests that multiple coordinated pathways converge to establish a permissive developmental environment in early postnatal life.

This bidirectional neuro-immune axis likely represents an adaptive physiological mechanism by which enteric circuitry is dynamically tuned to environmental conditions encountered after birth. By coupling neuronal development to immune signals that inherently respond to microbial colonization and dietary antigens, the host ensures that ENS maturation is calibrated to the specific postnatal niche. In this context, neonatal type 2 cytokine signaling reflects a transient developmental state optimized for tissue growth and remodeling rather than classical immune defense (15, 32).

Our study provides a conceptual framework for how immune-derived signals are integrated into the developmental programs of the human ENS, with direct relevance to pediatric gastrointestinal disorders such as necrotizing enterocolitis and Hirschsprung disease, where neuro-immune homeostasis is fundamentally disrupted. Future work incorporating spatially resolved transcriptomics and in vivo models will be important to confirm the physical dynamics of neuron-T cell interactions and to extend these findings across the full complexity of the intestinal microenvironment.

## Materials and Methods

### Experimental Design

For this study we investigated bidirectional neuro-immune interactions by integrating human tissue transcriptomics, induced pluripotent stem cell–derived enteric neural lineages, and large-scale computational interactome analyses. All experiments were conducted with Ethical Approval 23-457-B of the local Ethics Committee (Friedrich-Alexander-Universität Erlangen-Nürnberg and Human Research Ethics committees of the Royal Victorian Eye and Ear Hospital (11/1031H, 13/1151H-004), University of Melbourne (1545394), University of Tasmania (H0014124) and the University of Western Australia (RA/4/1/5255) as per the requirements of the National Health and Medical Research Council of Australia (NHMRC)). iPSC lines were obtained from healthy control individuals at the Universitätsklinikum Erlangen upon informed consent (22-289-Bp) and were performed in accordance with categories 1b and 2b of the International Society for Stem Cell Research (ISSCR) guidelines. All experimental work was conducted in accordance with the Declarations of Helsinki.

### Neonatal Ileum Acquisition and Processing

All human procedures were previously ethical approved by the local Ethics Committee of the Friedrich-Alexander-Universität Erlangen-Nürnberg (ID 23-457-B). Ileal surgery resections from neonates were obtained at the Pediatric Surgery of the University Hospital Erlangen and adult resections were collected at the Department of General and Visceral Surgery of the University Hospital Erlangen. Ileal resections were washed with cold PBS and cut into 300-500mg pieces. Tissue was frozen in 1 ml CryoStor® CS10 (STEM-CELL Technologies) in a freezing container and stored at - 80°C until further processing. For RNA isolation, tissue embedding and cell isolation tissue was thawed as previously described (33).

### RNA Isolation from tissue and iPSC

Approximately 50 mg of tissue or cell pellet was placed in 1 ml TRIzol (Invitrogen) and homogenized following the medium soft protocol (6500rpm for 10s followed by 10 seconds pause for three cycles) in the Precellys 24 Touch Homogenizer (Bertin Instruments). The TRIzol homogenate was transferred to a new tube and 200 µl chloroform was added. This was also done in case of RNA isolation from cells which were directly lysed in TRIzol. The suspension was vigorously mixed for 15s before centrifugation at 18000xg speed for 15 min at 4°C. The clear upper phase was then combined with 70% ethanol at a 1:1 ratio. From here onwards the Qiagen RNeasy Mini Kit (Cat74104) was used according to manufacturer’s instructions including gDNA removal. RNA quantification was performed using the NanoPhotometer NP80 (Implen).

### Bulk RNASeq and Data Analysis

RNA quality was assessed prior to mRNA library preparation and sequencing, both performed by Novogene (Munich, Germany). Sequencing was carried out on an Illumina NovaSeq 6000 platform using paired-end analysis. Preprocessed reads were mapped to the Homo sapiens reference genome GRCh38. Briefly, adapter sequences were trimmed from raw FASTQ files (34), and the resulting reads were aligned to the reference genome using HISAT2 (v2.2.1) (35). A count matrix was subsequently generated with featureCounts (36). Differential expression analysis was performed using DESeq2 (v1.42.0) (37), applying a significance threshold of adjusted *p ≤* 0.05 and an absolute *log*2 *foldchange ≥*1. Gene set enrichment analysis (GSEA) was conducted using GSEA software (v4.3.2) (38) and clusterProfiler (v4.10.1).

### Immunofluorescence

Ileal resections were embedded in paraffin and cut at 3 µm for immunofluorescence staining. Slices were dewaxed at 60°C for 45 minutes and incubated three times in Roti Histol solutions for 5 minutes each, followed by 5-minute incubation in ethanol 100%, 96% and 70% consecutively. Samples were treated with fixative solution consisting of methanol, hydrogen peroxide and deionized water in a ratio of 100:3:23 (v/v/v) for 20 minutes at RT. Precisely, the solution was prepared by mixing 50mL of methanol, 1,5mL of 30% Hydrogen Peroxide and 11,5mL of deionized water. Samples were washed thrice with TBS and antigen retrieval was performed by microwaving samples at 420watt with Tris-EDTA Buffer for 20 minutes. Subsequently, samples were cooled were permeabilized in Triton X-100 (0.1%) for 10 minutes. Samples were washed thrice with TBS and blocked with 10% Roti-ImmunoBlock (Art-Nr. T144.1) diluted in BSA 2% for 15 minutes followed by another round of washings and one hour incubation with either goat or donkey serum 10%. Primaries were incubated in the dilutions according to the table (Supplementary Table 1) overnight at 4°C. Lastly, samples were washed three times and incubated with appropriate secondaries for one hour at RT followed by 10-minute Hoechst staining. Mounting was carried on Mowiol (Carl ROTH – Cat0713.2) and image acquisition was performed at the microscope Zeiss AxioCam ERc 5s.

### Image Analysis

Confocal images were processed and quantified using the Fiji distribution of ImageJ (39). To ensure spatial accuracy, all images were calibrated to physical units using metadata-derived scaling factors prior to analysis. HuC/D+ images were duplicated and converted to 8-bit grayscale format to optimize thresholding efficiency. Segmentation of HuC/D+ neuronal cell bodies were performed by applying a global (Otsu) auto-thresholding arbitrated by manual adjustment to minimize background noise. To resolve individual neurons within dense clusters or ganglia, a watershed binary segmentation algorithm was applied to the resulting masks. Automated quantification was conducted using the ‘Analyze Particles’ plugin. Particles were filtered by size (50–250 micron2) and circularity (0.20–1.00) to exclude non-neuronal debris and imaging artifacts. The total neuron count was normalized to the total area analyzed (mm2).

### Flow Cytometry

Cryopreserved tissue was thawed as described above, washed in ice-cold PBS and incubated in 5 mL of digestion medium (DMEM F12 Gibco, containing 10 mM HEPES, 200 µg/mL of DNase I (Cat. 11284932001, Merck), 3 mg/mL of Collagenase II (Cat. 17105-015, Gibco), 2.5mg/mL of Collagenase D (Cat. 11088866001, Roche) and 0.25 mg/mL of dispase (Cat. 17105-041, Gibco) in the gentleMACS C Tube (Cat. 130-093-237, Miltenyi Biotec). Subsequently, the C-tubes were placed in the gentleMACS Octo Dissociator for 1 h at 37°C following the manufacturer’s protocol.Tissue was homogenized by gentle up and down pipetting followed by dissociation by a 19G needle and filtering through a 70 µm cell strainer. 5 mL of PBS without Ca2+ and Mg2+ with 2% Fetal Calf Serum was added to the suspension and cells were centrifuged for 3 min at 550xg and resuspended in red blood cell lysis solution (ACK Buffer). Cells were centrifuged again at 550xg for 3 min, ressuspendend in PBS and counted with acridine orange in the Luna FL cell counter (Logos Biosystems). To detect intracellular cytokines, cells were incubated with both Cell Stimulation Cocktail (Cat. 00-4970-93, ThermoFisher Scientific) and Protein-Transport-Inhibitor-Cocktail (Cat. 00-4980-93, ThermoFisher Scientific) in DMEM/F12 medium at 37°C for 4 h. All antibodies we diluted according to the table (Supplementary Table 1). Firstly, cells were stained with viable marker (Zombie NIR) for 20 min at room temperature (RT) followed by two washing steps with PBS and then further stained with extra-cellular markers for 30 min at RT. Cells were then fixed in BD Fixation/Permeabilization Solution (Cat. 51-209KZ, BD Biosciences) for 15 min at RT, washed in PBS and permeabilized with BD Perm/wash Buffer (Cat. 51-2091KZ, BD Biosciences) at 4°C in the dark for 45 min. For intracellular markers cells were resuspended in Perm/Wash containing antibodies for 2 h at 4°C in the dark. Cell analysis was performed using the Spectral Viewer ID7000 (Sony) and acquired data was analyzed FlowJo 10.1 (BD Biosciences).

### t-SNE Analysis

To characterize the architectural diversity of the immune landscape, we performed t-SNE manifold learning. To ensure an unbiased comparative analysis across disparate lineages, we normalized the input data using a downsampling algorithm to achieve equivalent event counts for T cells (CD45+ CD3+ CD8- CD19-), Innate Lymphoid Cells (ILCs) (CD45+ CD3- CD8- CD19- CD177- CD68-), and macrophages (CD45+ CD3- CD8- CD19- CD177- CD68+). High-dimensional clustering was executed on individual samples using a panel of lineage-defining markers and IL-13.

By projecting IL-13 expression density onto the resulting t-SNE coordinates, we identified specific cellular clusters associated with elevated cytokine production. These populations were subsequently quantified via manual gating to determine the total frequency of IL-13+ events relative to the total immune compartment in each sample.

### iPSCs Culture and Enteric Neuronal Lineage Differentiation

We employed six healthy control lines of which fibroblasts were obtained from patients (22-289-Bp) at the Universitätsklinikum Erlangen or University of Melbourne. iPSC vials were thawn in 37°C water bath, spun down and resuspended in Essential 8 Flex Media (Thermo Fisher) with 1% of Penicillin-Streptomycin and Rock Inhibitor (10 µM) for 24 h. Expansion and passaging were performed using the same media without Rock Inhibitor until required cell number was reached for differentiation. Commitment to enteric neuronal phenotype was provided as previously described (21). On day 0, confluent iPSC (70-80%) were incubated with Cocktail A (Essential 6 medium (Thermo Fisher), 10 µM SB431542 (Cat130-106-543, Miltenyi Biotec)), 600 nM CHIR99021 (Cat4423, Tocris), 1 ng/ml BMP4 (Cat120-05ET, Peprotech), 1% Penicillin/Streptomycin) for two days, followed by Cocktail B (Essential 6 medium (Thermo Fisher), 10 µM SB431542, 1.5 µM CHIR99021, 1% Penicillin/Streptomycin) replacement every two days for four days. Next, Cocktail C (Essential 6 medium (Thermo Fisher), 10 µM SB431542, 1.5 µM CHIR99021, 1 µM RA, (CatR2625, Sigma) 1% Penicillin/Streptomycin) was added and replaced every two days for six days. After Cocktail C, cells were detached with accutase (Thermo Fisher) and cultured in ultra-low attachment plates with Neurobasal medium (Thermo Fisher) containing 1x N2 supplement (Cat17502048, ThermoFisher Scientific), 1x B27 supplement w/o vitamin A (Cat11140050, ThermoFisher Scientific), 1x GlutaMax (Gibco – Cat35050038), 1x MEM NEAA (Gibco – Cat1114005), 10 ng/ml FGF2 (Peprotech – Cat100-18B), 3 µM CHIR99021, 1% Penicillin/Streptomycin) for three days to form spheroids. Finally, spheroids were digested with accutase and single cell neuron progenitors were plated and kept in Neurobasal medium (Thermo Fisher), 1x N2 supplement, 1x B27 supplement with Vit. A (Cat17504044, ThermoFisher Scientific), 1x GlutaMax, 1x MEM NEAA, 100 µM Ascorbic Acid (CatA4544, Sigma), 10 ng/ml GDNF (Cat450-10, Peprotech), 1% Penicillin/Streptomycin until day 70.

### Calcium Imaging

Live-cell Ca2+ imaging was performed on differentiated neurons treated with IL-13 (25 or 50 ng/mL) or vehicle control (PBS) at d21 and d30 of differentiation across three iPSC lines. Briefly, cells were incubated with 5 µM Fluo-4 AM (Invitrogen) in HEPES-buffered solution (149 mM NaCl, 5 mM KCl, 1 mM MgCl2, 2 mM CaCl2, 10 mM glucose, and 10 mM HEPES; pH 7.4) for 20 min at room temperature, followed by a 20 min wash in fresh HEPES solution. Dishes were then transferred to a recording chamber and continuously perfused with fresh HEPES solution maintained at 37°C. Recordings were acquired at 2 Hz with 50 ms exposure times for 1–2 min using an inverted fluorescence microscope (Axiovert 25, Zeiss) equipped with a 20× objective. Fluo-4 AM was excited at 470 nm with an LED (Zeiss Colibri) and its fluorescence emission was collected at 525 nm Recordings included a 5 s baseline period followed by local application of agonists, including high potassium solution (5 s; 78 NaCl, 75 KCl, 10 HEPES, 2 CaCl2, 1 MgCl2, and 10 D-glucose) and 1,1-dimethyl-4-phenylpiperazinium (DMPP; 10 s). Videos were analysed in IGOR PRO (Wavemetrics, Lake Oswego, OR, USA) using custom-written macros from Prof Pieter Vanden Berghe (KU Leuven, Belgium) (40). Regions of interest were manually drawn around individual cell bodies. Fluorescence intensity was normalized to baseline fluorescence (F0), calculated as the average signal during the first 10 frames, and expressed as Δ*F*/F0. Only cells exhibiting a minimum 5-fold increase above baseline noise were included in the analysis. Peak amplitude was defined as the maximal Δ*F*/F0 response. Statistical analyses were performed using Tukey’s multiple comparisons test in GraphPad Prism v9.

### scRNA-seq

#### Filtering and quality controls

Pre-processed single-cell RNA sequencing data was obtained from the Pan-Gastrointestinal (Pan-GI) Gut Cell Atlas (41). As described in the original dataset generation, tissue samples were dissociated and processed using the 10x Genomics Chromium platform, with upstream computational pre-processing. Quality control and original batch-effect integration across multi-donor fetal, pediatric, and adult cohorts were retained from the original Pan-GI Seurat object prior to targeted downstream analysis. All downstream data processing, filtering, and visualization were performed in the R statistical programming environment using the Seurat package (v5.0) and SingleCellExperiment, with file formatting and storage managed by the SeuratDisk package. To focus the analysis on targeted small intestine cell populations, the overarching healthy and diseased Pan-GI atlas was computationally subsetted based on clinical and spatial metadata parameters. Cells were restricted exclusively to the small intestine by filtering for annotations specifying the duodenum, jejunum, ileum, and general small intestine. The cohort was further filtered to encompass healthy baseline controls (annotated as control and organ donor) alongside specific clinical states, including intestinal atresia, Familial Adenomatous Polyposis, and fistula revisions. Subsequently, cells were stratified into two distinct developmental groups based on donor age: a Neonate group comprising first-trimester (6-13 weeks), second-trimester (14-20 weeks), and preterm (23-31 weeks) samples, and an adult group encompassing donors spanning 18 to 80 years of age.

#### Compositional analysis

For comparative analysis, subsets were manipulated as H5Seurat objects, and two-dimensional visualizations were generated using Uniform Manifold Approximation and Projection (UMAP) embeddings derived from the integrated Pan-GI spatial mapping. Cellular populations were evaluated at two hierarchical annotation resolutions and split by developmental stage to allow for localized developmental comparisons. Compositional analysis was performed using the scProportionTest package (v0.0.0.9000) (42), starting with the metadata containing the cell types and developmental stage information. The sc utils function was applied to process the metadata, and permutation testing was then conducted to evaluate whether the proportions of cells in specific clusters differed significantly between the two groups (Neonates vs. Adults). The analysis was run with a false discovery rate (FDR) threshold of 0.05 and a *log*2*foldchange ≥* 1.

#### Inference of Signaling Networks

To characterize the intercellular signaling landscapes of the small intestine, we performed cell-cell communication analysis using the CellChat (v1.6.1) R package (23). The Seurat object was partitioned into two distinct datasets based on developmental stage: Neonates and Adults. For each cohort, a CellChat object was initialized using the human ligand-receptor database (CellChatDB.human), encompassing over 2,000 documented interactions. Data was pre-processed by identifying over-expressed genes and interactions within each group.

#### Communication Probability and Pathway Analysis

The communication probability for each ligand-receptor pair between a sender cell group and a receiver cell group was calculated using a mass action law-based model, incorporating a minimum cell threshold of 10 cells per group to ensure robust inference. For each developmental cohort, we computed the communication probability at the level of individual ligand-receptor pairs and summarized them into coordinated signaling pathways. Centrality measures, including out-degree (sender), in-degree (receiver), mediator, and influencer scores, were calculated to identify key orchestrators of signaling within each network.

#### Comparative Signaling Analysis

To identify developmental shifts in signaling, we performed a comparative analysis between the Neonate and Adult objects. Total interaction counts and overall interaction strengths were compared to quantify the global signaling flux. We utilized hierarchical heatmap visualizations and dot plots to identify upregulated and downregulated pathways across the different cell lineages.

#### Visualization of Targeted Interactions

Cell-state-specific communication was visualized using chord diagrams to map directional signaling between Neural and T and NK cell populations. Differential signaling roles for each cell type were further characterized using network centrality plots, highlighting the relative importance of each lineage as a sender or receiver. For high-resolution analysis of specific pathways, we generated violin plots to assess the expression levels of ligands and their cognate receptors across all major lineages.

#### Characterization of IL-13 signaling

To characterize the expression of specific signaling components, average expression profiles for genes within the IL13 pathway were calculated across each lineage and developmental stage. Row-scaled Z-scores were visualized using a heatmap generated via ComplexHeatmap, with columns split by cell type and annotated by developmental condition. To quantify differences in specific downstream mediators, expression levels were extracted from the Neural and T and NK cell subsets. Statistical significance between Neonate and Adult groups was determined using the Wilcoxon rank-sum test with Benjamini-Hochberg p-value adjustment. These distributions were visualized using violin plots incorporating internal boxplots to illustrate median and interquartile ranges.

### Statistical Analysis

All results were firstly tested for normality distribution. Samples under a normal distribution, verified by Shapiro-Wilk test, were submitted to unpaired two-tailed student t-test. For samples acquired and compared between different time-points, Two-way ANOVA testing was used with Tukey’s multiple comparisons test. For correlation analysis samples were tested by the spearman test. For group comparison of sequenced samples between neonates and adults Wilcoxon Sum Rank t-test was employed. Samples were considered statistically significant when p < 0.05. Statistical analysis was performed using GraphPad Prism 10.

## Data availability

RNA sequencing data have been deposited at Zenodo (10.5281/zenodo.19567405) and are publicly available as of the date of publication. Other data and images that support the findings of this study are available on request from the lead contact. Any additional information required to reanalyze the data reported in this paper is available from the lead contact upon request.

## ACKNOWLEDGEMENTS

We thank the participating neonates and families, the adult tissue donors and the generous and inspirational support of Anton’s parents. Moreover, we thank Daniel Beß and Katharina Rost for their generous support to the project. Additionally, the authors would like to thank Professor Alice Pebay for provision of some of the iPSC lines used in this study (43), and acknowledge the facilities and the scientific and technical assistance of the iPSC reprogramming facilities, the Stem Cell Disease Modelling laboratory, the University of Melbourne, which is supported by Phenomics Australia (PA) through funding from the Australian Government’s National Collaborative Research Infrastructure Strategy program. This research has received funding from the Deutsche Forschungsgemeinschaft (DFG [German Research Foundation]) – SFB/TRR369 DIONE – 501752319 (A02). Further support was provided by the Interdisciplinary Center for Clinical Research of the Friedrich-Alexander-Universität Erlangen-Nürnberg (Pilot project funding P175, Junior project funding J117 and P169 (ELAN funding 24-07-28-1-Diez (SD)). MN and BW were Funded by the Deutsche Forschungsgemeinschaft (DFG, German Research Foundation) – 505539112 – KFO 5024 (A01). Further support came from the Bavarian Ministry of Science and the Arts in the framework of the ForInter network. S.P.R. was supported by the Deutsche Forschungsgemeinschaft (DFG, German Research Foundation), Emmy Noether Programme RO 6247/1-1 (project ID 446316360), DFG SFB1160 (project ID 256073931), SFB 1755 (project ID 550296805) TRR 359 (project ID 491676693), TRR 417 (project ID 540805631).

## AUTHOR CONTRIBUTIONS

Conceptualization: LEDG, BG, IVH and SD; Methodology: LEDG, BG, AP, HY, LR, OT, AN, AJ, LT, MW, SPR and IVH; Investigation: LEDG, BG, MMH, LAS, IVH and SD; Visualization: LEDG, BG, LR, AN, MMH, LAS, IVH and SD; Supervision: MB, MN, CG, BW, IVH and SD; Biomaterial: CS and ME; Grant Acquisition and Resources: MN, CG, BW, IVH and SD; Writing-original draft: LEDG, BG, AN, MMH, LAS, IVH and SD; Writing-review and editing: LEDG, BG, IVH and SD

## Supplementary Figures

**Supplementary figure 1.**
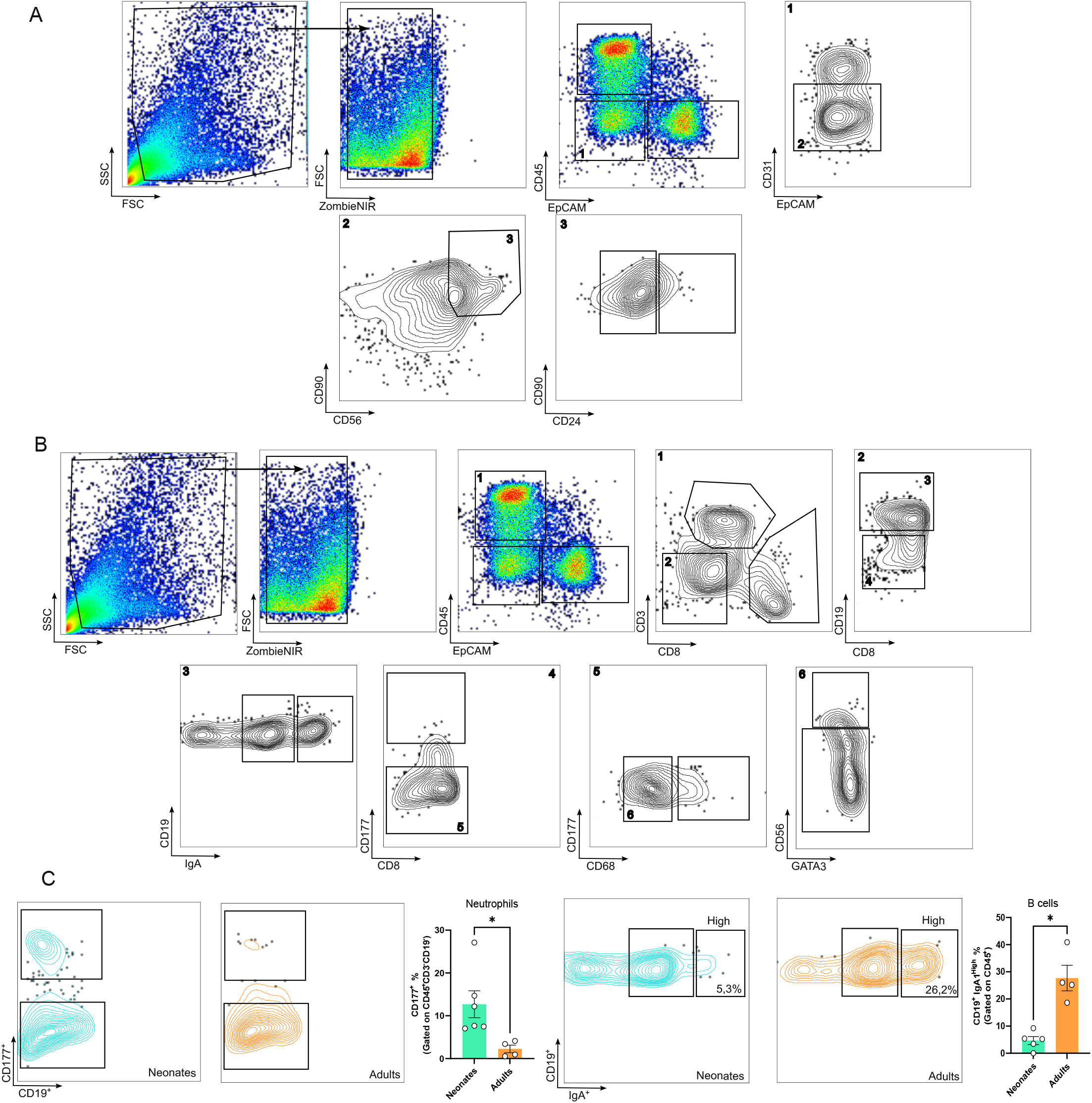
(A) Gating panel of enteric neuronal and glial cells separation by flow cytometry where neurons are CD45-CD31-CD90+CD56+CD24high and glial cells are CD45-CD31-CD90+CD56+CD24+. (B) Gating strategy to visualize immune cells in the ileum. T cells are gated either as CD45+CD8+ or CD45+CD3+; B cells are gated as CD45+CD19+CD3- and/or IgA+. Neutrophils are gated as CD45+CD3-CD19-CD177+ and macrophages as CD45+CD3-CD19-CD177-CD68+. (C) Neutrophil and IgA+ B cells gating strategy along with frequencies in neonates and adults.

**Supplementary figure 2.**
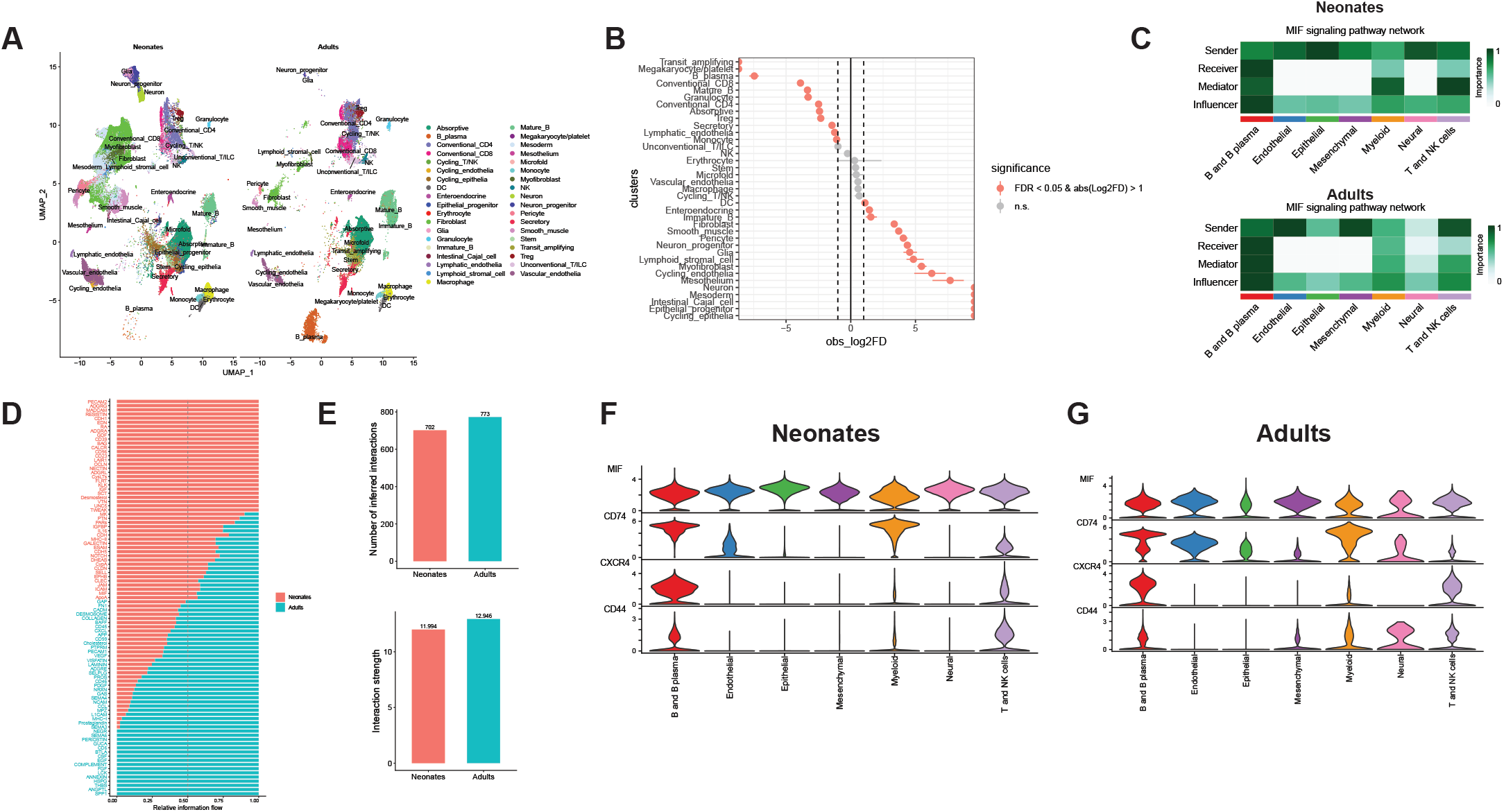
(A) UMAP visualization of the global cellular landscape, color-coded by major cell lineages with a higher level of annotation. (B) Compositional analysis using scProportion test. Significant differences are defined by FDR < 0.05 and abs(Log2FD) > 1. (C) Hierarchical plot showing the MIF signaling pathway network in neonates and adults. Color scale denotes the importance, distinguishing roles between Sender, Receiver, Mediator, and Influencer. (D) Relative information flow across signaling pathways. Pathways enriched in neonates are highlighted in pink, while those enriched in adults are highlighted in green. (E) Bar charts comparing the total number of inferred interactions and overall interaction strength between the neonatal and adult datasets. (F-G) Violin Plots showing expression of the MIF pathway components in the cell clusters, specifically highlighting interactions involving MIF and its receptors CXCR4, CD44, and CD74 in neonates (F) and adults (G).

## Bibliography

1. M. C. Arrieta, L. T. Stiemsma, N. Amenyogbe, E. M. Brown, B. Finlay, The intestinal microbiome in early life: health and disease. Frontiers in immunology 5, 427 (2014).

2. J. B. Furness, The enteric nervous system and neurogastroenterology. Nature Reviews Gastroenterology and Hepatology 9, 286–294 (2012).

3. G. O. Adeniyi-Ipadeola et al., Infant and adult human intestinal enteroids are morphologically and functionally distinct. mBio 15, e01316–01324 (2024).

4. L. C. Frazer, M. Good, Intestinal epithelium in early life. Mucosal Immunology 15, 1181–1187 (2022).

5. E. Drokhlyansky et al., The Human and Mouse Enteric Nervous System at Single-Cell Resolution. Cell 182, 1606–1622.e1623 (2020).

6. H. B. Mikkelsen, L. Thuneberg, Op/op mice defective in production of functional colony-stimulating factor-1 lack macrophages in muscularis externa of the small intestine. Cell and Tissue Research 295, 485–493 (1999).

7. M. Stakenborg et al., Enteric glial cells favor accumulation of anti-inflammatory macrophages during the resolution of muscularis inflammation. Mucosal Immunology 15, 1296–1308 (2022).

8. V. Grubišić et al., Enteric Glia Modulate Macrophage Phenotype and Visceral Sensitivity following Inflammation. Cell Reports 32, 108100 (2020).

9. A. Jarret et al., Enteric Nervous System-Derived IL-18 Orchestrates Mucosal Barrier Immunity. Cell 180, 50–63.e12 (2020).

10. L. A. Schwerdtfeger, S. A. Tobet, Vasoactive intestinal peptide regulates ileal goblet cell production in mice. Physiological Reports 8, e14363 (2020).

11. M. Herath, S. Hosie, J. C. Bornstein, A. E. Franks, E. L. Hill-Yardin, The Role of the Gastrointestinal Mucus System in Intestinal Homeostasis: Implications for Neurological Disorders. Frontiers in Cellular and Infection Microbiology Volume 10-2020, (2020).

12. G. M. H. Birchenough, M. E. Johansson, J. K. Gustafsson, J. H. Bergström, G. C. Hansson, New developments in goblet cell mucus secretion and function. Mucosal Immunology 8, 712–719 (2015).

13. A. Dariel et al., Analysis of enteric nervous system and intestinal epithelial barrier to predict complications in Hirschsprung’s disease. Scientific Reports 10, 21725 (2020).

14. J. R. Thiagarajah et al., Altered Goblet Cell Differentiation and Surface Mucus Properties in Hirschsprung Disease. PLOS ONE 9, e99944 (2014).

15. R. M. Barilla et al., Type 2 cytokines act on enteric sensory neurons to regulate neuropeptide-driven host defense. Science 389, 260–267 (2025).

16. A. Olin et al., Stereotypic Immune System Development in Newborn Children. Cell 174, 1277–1292.e1214 (2018).

17. J. D. Windster et al., A combinatorial panel for flow cytometry-based isolation of enteric nervous system cells from human intestine. EMBO Reports 24, EMBR202255789 (2023).

18. S. Li et al., Interleukin-13 and its receptor are synaptic proteins involved in plasticity and neuroprotection. Nat Commun 14, 200 (2023).

19. H. Wang, J. P. P. Foong, N. L. Harris, J. C. Bornstein, Enteric neuroimmune interactions coordinate intestinal responses in health and disease. Mucosal Immunology 15, 27–39 (2022).

20. J. J. Barron et al., Group 2 innate lymphoid cells promote inhibitory synapse development and social behavior. Science 386, eadi1025 (2024).

21. K. Barber, L. Studer, F. Fattahi, Derivation of enteric neuron lineages from human pluripotent stem cells. Nature Protocols 14, 1261–1279 (2019).

22. A. J. Oliver et al., Single-cell integration reveals metaplasia in inflammatory gut diseases. Nature 635, 699–707 (2024).

23. S. Jin et al., Inference and analysis of cell-cell communication using CellChat. Nature Communications 12, 1088 (2021).

24. M. Ota et al., CD23+IgG1+ memory B cells are poised to switch to pathogenic IgE production in food allergy. Science Translational Medicine 16, eadi0673 (2024).

25. T. J. R. Frith et al., Retinoic Acid Accelerates the Specification of Enteric Neural Progenitors from <em>In-Vitro</em>-Derived Neural Crest. Stem Cell Reports 15, 557–565 (2020).

26. J. D. Eisenberg et al., Three-Dimensional Imaging of the Enteric Nervous System in Human Pediatric Colon Reveals New Features of Hirschsprung’s Disease. Gastroenterology 167, 547–559 (2024).

27. R. Hamnett et al., Regional cytoarchitecture of the adult and developing mouse enteric nervous system. Current Biology 32, 4483–4492.e4485 (2022).

28. P. Parathan, Y. Wang, A. J. L. Leembruggen, J. C. Bornstein, J. P. P. Foong, The enteric nervous system undergoes significant chemical and synaptic maturation during adolescence in mice. Developmental Biology 458, 75–87 (2020).

29. L. B. Dershowitz, J. A. Kaltschmidt, Enteric Nervous System Striped Patterning and Disease: Unexplored Pathophysiology. Cellular and Molecular Gastroenterology and Hepatology 18, 101332 (2024).

30. L. B. Dershowitz, L. Li, A. M. Pasca, J. A. Kaltschmidt, Anatomical and functional maturation of the mid-gestation human enteric nervous system. Nature Communications 14, 2680 (2023).

31. E. M. Schill, A. N. Floyd, R. D. Newberry, Neonatal development of intestinal neuroimmune interactions. Trends in Neurosciences 45, 928–941 (2022).

32. Y. Wang et al., Bi-directional communication between intrinsic enteric neurons and ILC2s inhibits host defense against helminth infection. Immunity 58, 465–480.e468 (2025).

33. E. Gamero-Estevez et al., Protocol for Immune Cell Isolation, Organoid Generation, and Co-culture Establishment from Cryopreserved Whole Human Intestine. Bio Protoc 15, e5157 (2025).

34. L. Yan et al., Single-cell RNA-Seq profiling of human preimplantation embryos and embryonic stem cells. Nat Struct Mol Biol 20, 1131–1139 (2013).

35. A. Mortazavi,B. A. Williams, K. McCue, L. Schaeffer, B. Wold, Mapping and quantifying mammalian transcriptomes by RNA-Seq. Nat Methods 5, 621–628 (2008).

36. Y. Liao, G. K. Smyth, W. Shi, featureCounts: an efficient general purpose program for assigning sequence reads to genomic features. Bioinformatics 30, 923–930 (2014).

37. M. I. Love, W. Huber, S. Anders, Moderated estimation of fold change and dispersion for RNA-seq data with DESeq2. Genome Biol 15, 550 (2014).

38. A. Liberzon et al., The Molecular Signatures Database (MSigDB) hallmark gene set collection. Cell Syst 1, 417–425 (2015).

39. J. Schindelin et al., Fiji: an open-source platform for biological-image analysis. Nature Methods 9, 676–682 (2012).

40. Z. Li et al., Regional complexity in enteric neuron wiring reflects diversity of motility patterns in the mouse large intestine. Elife 8, (2019).

41. R. Elmentaite et al., Cells of the human intestinal tract mapped across space and time. Nature 597, 250–255 (2021).

42. S. A. Miller et al., LSD1 and Aberrant DNA Methylation Mediate Persistence of Enteroendocrine Progenitors That Support BRAF-Mutant Colorectal Cancer. Cancer research 81, 3791–3805 (2021).

43. M. Daniszewski et al., Retinal ganglion cell-specific genetic regulation in primary open-angle glaucoma. Cell Genom 2, 100142 (2022).

